# Equilibrium Propagation with Predictive Learning in Leaky Integrate-and-Fire Spiking Neural Networks

**DOI:** 10.64898/2026.05.19.726261

**Authors:** Yoshimasa Kubo

## Abstract

Equilibrium propagation (EP) is a biologically plausible alternative to backpropagation that has demonstrated competitive performance across a range of machine learning tasks. Recent work has extended EP to spiking neural networks (SNNs), leveraging leaky integrate-and-fire (LIF) neurons and spike-based plasticity rules to improve biological realism while maintaining strong performance. In this work, we propose an EP-based SNN framework that combines LIF neural dynamics with a predictive learning rule, replacing conventional spike-timing-dependent plasticity (STDP) with a learning rule more directly aligned with predictive coding principles. We evaluate the proposed model on multiple image classification benchmarks, including MNIST, KMNIST, and Fashion-MNIST, and compare its performance with a BP-trained LIF SNN baseline. Our results show that the proposed EP-based LIF model (EP+LIF) achieves competitive accuracy across datasets, with performance approaching that of the BP-trained counterpart (BP+LIF) while relying on a biologically motivated local learning rule. In addition, analysis of hidden-layer spiking activity reveals that EP+LIF produces more persistent hidden-state activity, whereas BP+LIF yields sparser spiking representations. These results demonstrate that predictive learning can support effective EP-based training in LIF spiking networks, while also highlighting differences in activity patterns that motivate future work on activity regulation and sparse spiking dynamics.

## Introduction

Equilibrium propagation (EP) is a biologically plausible learning algorithm and an alternative to backpropagation (BP) for training neural networks (Ernoult et al., 2019; Laborieux & Zenke, 2022; Laborieux et al., 2021; Scellier & Bengio, 2017, 2019). EP has been successfully applied to a variety of tasks, including image classification (Laborieux et al., 2021), continual learning (Kubo et al., 2025), and reinforcement learning (Kubo et al., 2022), where it achieves performance competitive with BP-based models. Recent work has also shown that incorporating biologically inspired neural mechanisms, such as dendritic neurons, can further improve the performance of EP-based models (Kubo, 2026). More recently, several studies have extended EP to spiking neural networks (SNNs) (Lin et al., 2024; Martin et al., 2021; O’Connor et al., 2019), demonstrating that biologically grounded implementations can achieve strong performance on benchmark tasks. For example, EqSpike (Martin et al., 2021) employs leaky integrate-and-fire (LIF) neurons to model neural dynamics and updates synaptic weights using spike-timing-dependent plasticity (STDP), achieving 97.6% test accuracy on MNIST with a fully connected architecture. These results suggest that EP provides a promising framework for training SNNs using biologically motivated dynamics and local learning rules.

In this study, we adopt LIF neurons for neural dynamics within the EP framework. Instead of using STDP for learning, we employ the predictive learning rule (Luczak et al., 2022) as an alternative biologically motivated update mechanism. This learning rule is local and is motivated by predictive coding principles, making it well suited for biologically plausible learning in SNNs. We evaluate the proposed EP-based LIF model (EP+LIF) on multiple image classification benchmarks, including MNIST, KMNIST, and Fashion-MNIST, and compare its performance with a BP-trained LIF SNN (BP+LIF) baseline. In addition to classification performance, we analyze hidden-layer spiking activity during the free phase to examine how EP-based and BP-based training shape internal spiking dynamics.

Our results show that the proposed EP+LIF model achieves competitive accuracy across datasets, demonstrating that predictive learning can support effective EP-based training in LIF spiking networks. The spike-activity analysis further reveals that EP+LIF produces more persistent hidden-layer activity than BP+LIF, whereas BP+LIF produces sparser spiking representations. This difference highlights that biologically motivated local learning and BP-based optimization can induce distinct internal activity patterns, and motivates future work on regulating spike sparsity and improving computational efficiency in EP-trained SNNs.

## Methods

### Model

We consider a fully connected network trained with equilibrium propagation (EP), where neural dynamics are implemented using leaky integrate-and-fire (LIF) neurons. The model consists of an input layer, a hidden layer, and an output layer with tied feedforward and feedback weights.

### Leaky Integrate-and-Fire Dynamics

Neural dynamics follow a discrete-time LIF model:

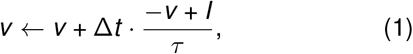

where *v* denotes the membrane potential, *I* the input current, *τ* the membrane time constant, and Δ*t* the integration step.

A spike *s* is emitted when *v* ≥ *v*_th_, after which the membrane potential is reset. To obtain stable activity for learning, we maintain a smoothed firing rate:

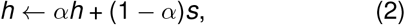

where *h* represents the smoothed neural activity and *α*∈ [0, 1] controls the rate smoothing.

### Equilibrium Propagation

The network evolves over multiple steps in two phases. In the *free phase*, the system converges with no target signal. In the *nudged phase*, the output layer is weakly driven toward the target:

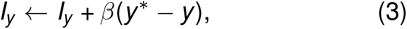

where *I*_*y*_ denotes the output input current and *β* controls the strength of the nudging signal.

Hidden and output inputs are given by:

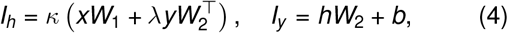

where *I*_*h*_ and *I*_*y*_ denote input currents to the hidden and output layers, respectively, *λ* is a feedback scaling parameter, and *κ* is an input scaling factor.

### Predictive Learning Rule

Weights are updated using differences between nudged-phase and free-phase activities:

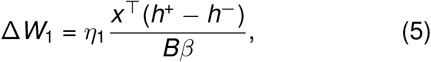

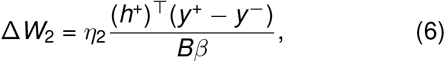

where *B* is the batch size and *β* is the nudging strength. Here, *h*^−^ and *y*^−^ denote the hidden and output activities at the end of the free phase, respectively, while *h*^+^ and *y* ^+^ denote the corresponding activities at the end of the nudged phase. The learning rates for the input-to-hidden and hidden-to-output weights are denoted by *η*_1_ and *η*_2_, respectively.

This update rule (Luczak et al., 2022) provides a local and biologically plausible alternative to backpropagation. In our implementation, we adopt the predictive learning rule without explicit prediction units (Kubo et al., 2023; Luczak & Kubo, 2022), resulting in a simplified biologically motivated formulation.

### Experimental Setup

Experiments were conducted on the MNIST (LeCun & Cortes, 2005), Kuzushiji-MNIST (KMNIST) (Clanuwat et al., 2018), and Fashion-MNIST (FMNIST) (Xiao et al., 2017) datasets. We used a fully connected architecture with a single hidden layer, and both the hidden and output layers employed leaky integrate-and-fire (LIF) neurons. For all the datasets, we evaluated hidden-layer sizes of 128, 256, 512, and 1024 neurons.

The main hyperparameters used for EP+LIF and BP+LIF are summarized in Table 1. For the BP+LIF baseline, we used the same fully connected LIF architecture and trained the model using backpropagation.

**Table 1:** Hyperparameters used for EP+LIF and BP+LIF experiments.

| Hyperparameter | EP+LIF | BP+LIF |
| --- | --- | --- |
| Epochs | 80 | 80 |
| Batch size | 64 | 64 |
| Hidden-layer size | 128, 256, 512, 1024 | 128, 256, 512, 1024 |
| Learning rate | $\eta_1 = 0.3, \eta_2 = 0.022$ | 0.3 |
| Free-phase / simulation time steps | 230 | 230 |
| Nudged-phase time steps | 60 | – |
| Nudging parameter | $\beta = 1.0$ | – |
| Feedback scaling | $\gamma = 1.0$ | – |
| Hidden threshold | 0.2 for MNIST/FMNIST; 0.1 for KMNIST | 0.2 |
| Output threshold | 0.5 for MNIST/FMNIST; 0.45 for KMNIST | 0.5 |
| Rate smoothing | $\alpha = 0.8$ | $\alpha = 0.8$ |
| Input scaling | $\kappa = 2.0$ for MNIST/KMNIST; $\kappa = 1.5$ for FMNIST | $\kappa = 2.0$ |
| Optimizer / update rule | local EP update | SGD without momentum |

We did not use momentum or adaptive optimizers such as Adam (Kingma & Ba, 2014) for BP+LIF.

Each model was trained from scratch six times on each dataset using different random seeds.

The implementation is available at: https://github.com/ykubo82/LIF_EP

## Results

Table 2 summarizes the test accuracies across all datasets. We also analyze the performance gaps between EP+LIF and BP+LIF across datasets in Figure 1. For MNIST and KMNIST, both EP+LIF and BP+LIF improved as the hidden-layer size increased, indicating that larger hidden representations enhanced classification performance. BP+LIF consistently achieved the highest accuracy, as expected for a model trained with backpropagation. However, EP+LIF remained competitive across all hidden-layer sizes. On MNIST, the performance gap between BP+LIF and EP+LIF increased modestly with hidden-layer size, but remained small, reaching only 0.50 percentage points at 1024 hidden neurons. On KMNIST, EP+LIF closely followed BP+LIF across hidden sizes, with the gap remaining below one percentage point for hidden sizes of 256, 512, and 1024. On FMNIST, EP+LIF also achieved competitive performance, although the gap with BP+LIF was larger and the performance gain from increasing hidden size was more modest. This trend suggests that the more challenging FMNIST benchmark may benefit more from BP-based optimization and is further discussed in the Discussion section.

**Table 2:**
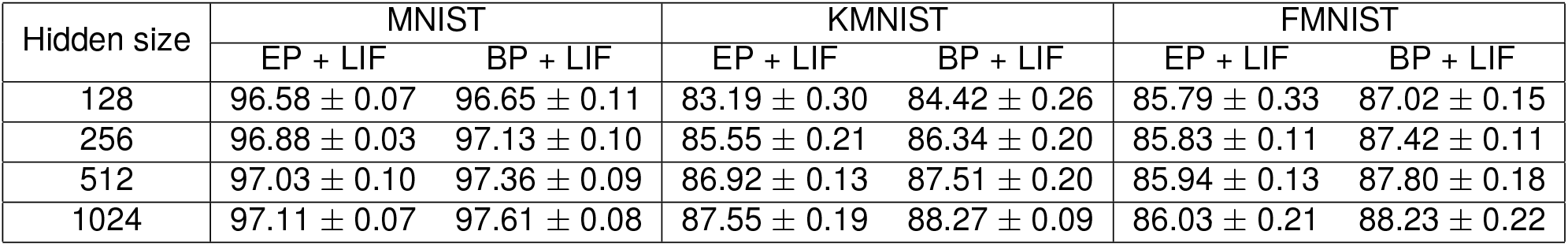
Average test accuracies (± standard deviation) of EP-trained and BP-trained LIF spiking neural networks on MNIST, KMNIST, and FMNIST.

**Figure 1:**
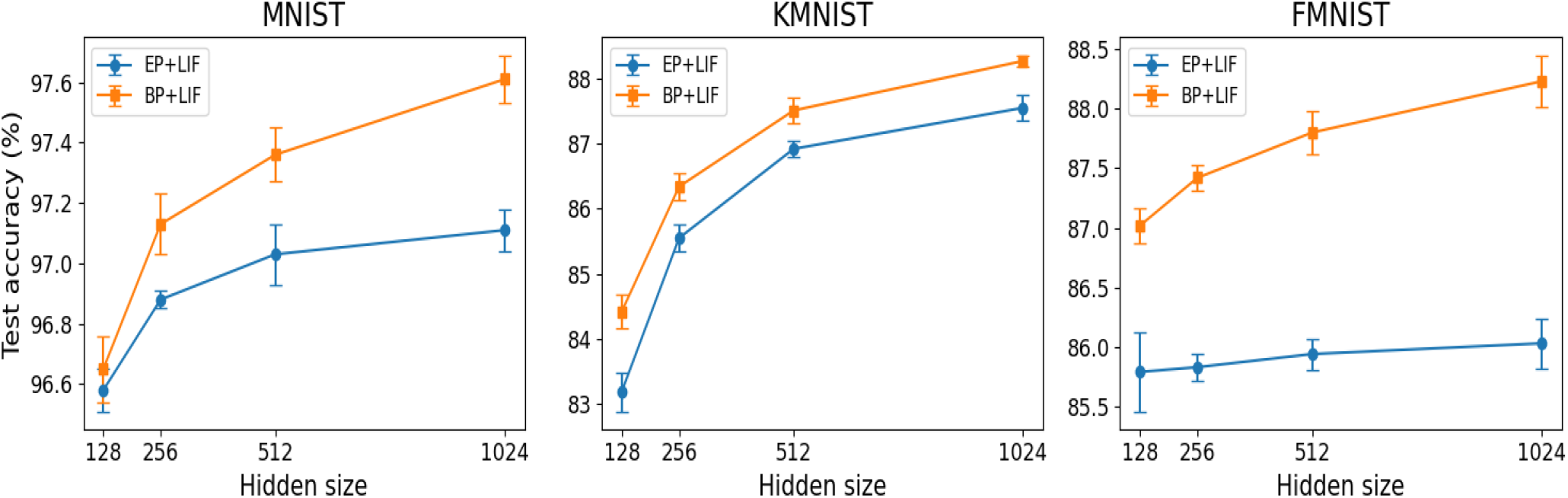
Accuracies for EP+LIF and BP+LIF vesus hidden sizes.

To further examine the learned spiking dynamics, we analyzed hidden-layer spike activity during the free phase (Figure 2) on the MNIST dataset. EP+LIF exhibited substantially higher average hidden-layer spike activity than BP+LIF, with a larger fraction of neurons showing persistent activity. Specifically, the mean hidden spike rate was 0.600 for EP+LIF and 0.175 for BP+LIF. In addition, 401 hidden neurons in EP+LIF had spike rates above 0.8, compared with only 7 neurons in BP+LIF. In contrast, BP+LIF produced a sparser distribution of hidden activity, with most neurons concentrated in the low-spike-rate regime. These results suggest that EP-trained LIF networks may rely on more sustained hidden-state activity, whereas BP-trained LIF networks produce sparser hidden representations.

**Figure 2:**
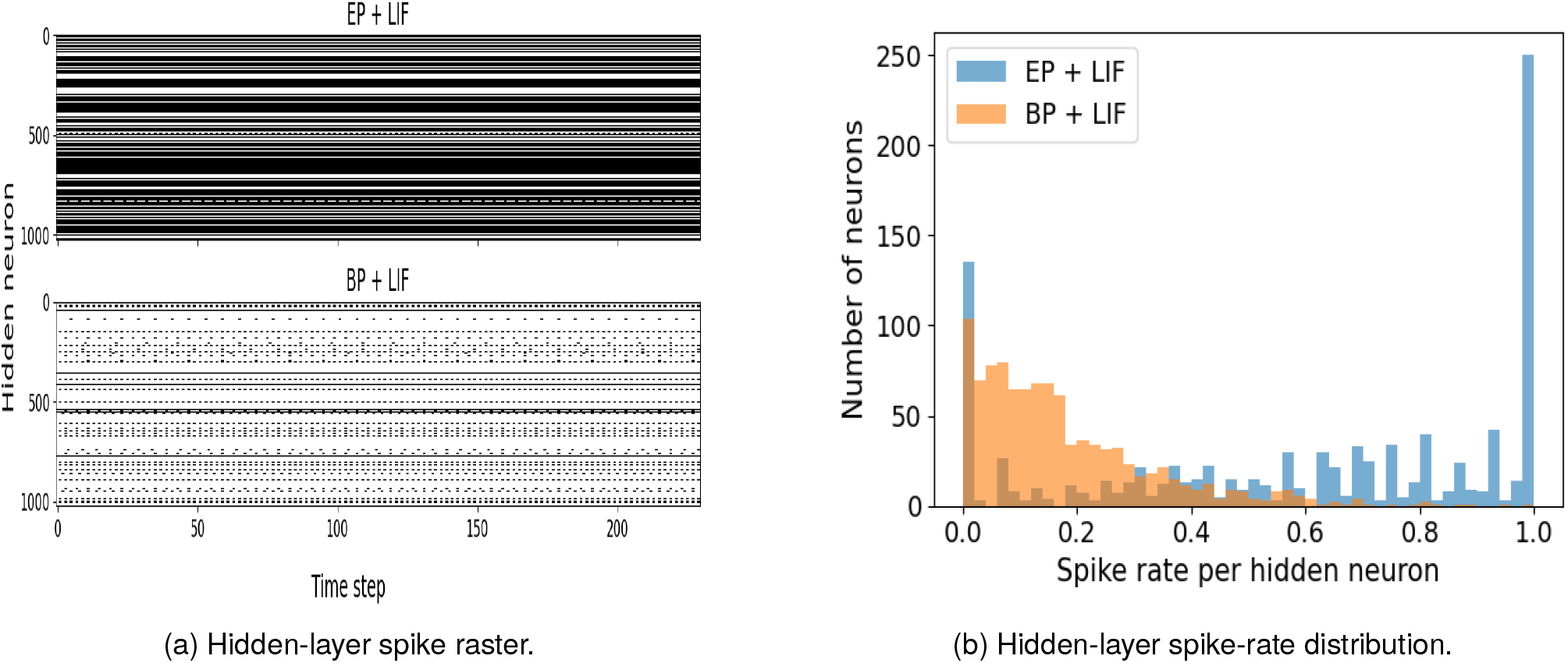
Hidden-layer spiking activity during the free phase for EP+LIF and BP+LIF models on MNIST. (a) Raster plots show that EP+LIF exhibits more persistent hidden-layer spiking activity, whereas BP+LIF produces sparser activity patterns. (b) The distribution of spike rates further shows that EP+LIF contains a larger fraction of persistently active hidden neurons, while BP+LIF is more concentrated in the low-spike-rate regime.

## Discussion

In this study, we introduced a spiking neural network (SNN) framework with leaky integrate-and-fire (LIF) neurons trained using equilibrium propagation (EP) and a predictive learning rule. We evaluated the proposed EP+LIF model on multiple image classification benchmarks, including MNIST, KMNIST, and Fashion-MNIST, and compared its performance with a BP-trained LIF SNN baseline across different hidden-layer sizes. The results showed that EP+LIF achieved competitive accuracy across datasets, demonstrating that predictive learning can support effective EP-based training in LIF spiking networks.

Our work is closely related to prior EP-based SNN studies such as EqSpike (Martin et al., 2021), which combined LIF neural dynamics with STDP-based synaptic updates and achieved strong performance on MNIST. In contrast, the present study replaces STDP with a predictive learning rule motivated by predictive coding principles. Although the architectures and training procedures are not identical, our EP+LIF model achieved 97.11% accuracy on MNIST with 1024 hidden neurons, which is close to the 97.6% MNIST accuracy reported by EqSpike using a fully connected architecture with 300 hidden neurons. While EqSpike achieved this result with a smaller hidden layer, the present study extends the evaluation beyond MNIST by testing the proposed model on KMNIST and Fashion-MNIST across multiple hidden-layer sizes. These results suggest that predictive learning provides an effective alternative local update mechanism for EP-trained spiking networks and can generalize across multiple image classification benchmarks.

Although BP+LIF achieved higher accuracy in most settings, this gap is expected because BP directly optimizes the network through gradient-based training. In contrast, EP+LIF relies on a biologically motivated local learning rule based on the difference between free and nudged phases. Importantly, the performance gap remained relatively small on MNIST and KMNIST, indicating that EP+LIF can approach BP-trained performance while avoiding direct backpropagation through the network. On Fashion-MNIST, the gap was larger across hidden-layer sizes; at 1024 hidden neurons, EP+LIF achieved 86.03% accuracy compared with 88.23% for BP+LIF. Interestingly, while BP+LIF showed consistent performance gains as the hidden-layer size increased from 128 to 1024, EP+LIF exhibited a more saturated trend on Fashion-MNIST. This suggests that simply increasing the capacity of a single hidden layer may not fully overcome the representational or optimization constraints imposed by local EP-based updates on more complex visual datasets. These results motivate future extensions toward richer architectures, such as convolutional or multi-layer models, as well as activity-regulating mechanisms that may improve learning efficiency in larger EP-trained SNNs. Nevertheless, the competitive performance across all three datasets supports the view that EP-based local learning can provide a promising biologically motivated alternative for training SNNs.

The spike-activity analysis further revealed a clear difference between EP+LIF and BP+LIF. EP+LIF produced more persistent hidden-layer activity, whereas BP+LIF produced sparser hidden representations. This difference indicates that biologically motivated local learning and BP-based optimization can induce distinct internal activity patterns, even when the same LIF architecture is used. In EP+LIF, sustained hidden activity may help maintain stable internal states during the free phase and support the contrastive update between free and nudged dynamics. However, higher spike rates may also reduce one of the practical advantages of SNNs, namely sparse event-driven computation and potential energy efficiency (Göltz et al., 2021; Oh et al., 2022). Moreover, sparse activity has long been discussed as an important coding principle in sensory systems (Olshausen & Field, 2004), suggesting that excessive persistent activity may also be undesirable from a biological perspective. Therefore, the persistent activity observed in EP+LIF should be interpreted as both a characteristic of the learned dynamics and a limitation that motivates further mechanisms for regulating spike sparsity.

One promising direction for regulating hidden-layer activity is the incorporation of heterogeneous time constants (Kubo et al., 2026; Perez-Nieves et al., 2021). Previous work on SNNs has shown that heterogeneous time constants can improve performance across multiple tasks (Perez-Nieves et al., 2021). In addition, prior work on non-spiking EP-based models has shown that heterogeneous temporal dynamics can improve learning stability and performance (Kubo et al., 2026). Extending these ideas to EP-trained LIF networks may help reduce excessive persistent activity while maintaining accuracy, thereby improving both learning dynamics and computational efficiency.

Another important direction is the use of dendritic neuron models, which have been studied as biologically grounded mechanisms for improving neural computation and credit assignment (Chavlis & Poirazi, 2025; Grewal et al., 2021; Kubo, 2026). Combining EP-based learning with dendritic architectures may provide a more expressive and biologically realistic framework for training SNNs. Such mechanisms may also help shape local activity patterns, regulate hidden-layer dynamics, and improve credit assignment without relying on standard backpropagation. Future work will investigate how heterogeneous time constants, dendritic computation, convolutional or multi-layer architectures, and other activity-regulating mechanisms can further improve the performance, stability, sparsity, and efficiency of biologically plausible learning in spiking neural networks.

## Acknowledgments

This research was enabled in part by computational resources provided by the Digital Research Alliance of Canada (alliancecan.ca).

## Notes

### Competing Interest Statement

The authors have declared no competing interest.

